# Cross-sectional and longitudinal research approaches: running pace and step characteristics of elite athletes in the 400-m hurdles

**DOI:** 10.1101/410464

**Authors:** Mitsuo Otsuka, Tadao Isaka

## Abstract

The aim of this study was to clarify the running pace and step characteristics among various competitive-level 400-m hurdlers through cross-sectional and longitudinal research approaches. We analysed spatiotemporal data for 13 male world-class and 14 male national-level 400-m hurdlers. We analysed 16.8 ± 4.2 races for each world-class hurdler and 20.0 ± 6.0 races for each national-level hurdler (the total number of analysed runs was 499) using publicly available television and internet broadcasts. Cross-sectional approach showed that positive relationships of finish time were highly obtained between both first- (r = 0.901) and latter-half split times (r = 0.914). In contrast, the first- and latter-half split times were not significantly correlated to SL and SF. A multiple single-subject approach showed that 14/27 hurdlers were identified as being latter-half speed reliant. In contrast, no hurdlers demonstrated first-half reliance. In the latter half of the race, 12/27 hurdlers were identified as being SF reliant; no hurdlers demonstrated SL reliance. In conclusions, important findings regarding high performance in a cross-sectional research approach do not always corresponded with those in a longitudinal research approach. Athletes and coaches should carefully improve performance in the first half of the race based on an individualization principle for training.

## Introduction

The 400-m hurdles is a long sprint running event. During the long sprint event, the anaerobic glycolytic system is largely converted to the mechanical energy [1–6], which can be externally and kinematically observed using a video camera [5,7]. In particular, researchers can easily analyse spatiotemporal parameters because they can calculate these parameters just by visually counting the number of steps in the fixed moving distance. Therefore, a number of samples can be obtained, for example, through public data analysis [8].

In a 400-m hurdles race for men, a total of ten hurdles at 91.4 cm height are positioned 35 m apart. Hurdlers are required to minimize decelerations into running direction during clearing hurdles and to maintain high running speed during all inter-hurdle distances. Split times are often used to evaluate the running speed cross-sectionally and longitudinally and these are measured during the hurdles race using the instant of touchdown for the leading leg [9]. Compared to split times determined by touchdown method, it seems to be more difficult during competitive races that split times are determined measured using video camera or photocell, which are the gold standard methods for the experimental measurements [10]. This is because that detecting the instant when the torso frontally passes one thin line is visually difficult using the panned video camera, or the photocell device cannot be directly set close to athletes.

Race pace is a key strategy for a good finish time in a long sprint running event [12]. A cross-sectional research reported that the faster the finish times of 400-m hurdlers are, the faster touchdown split-times are during the latter half of the race [11]. However, because “correlation does not imply causation,” it is unclear whether the first- or latter-half split time is more important for each elite hurdler to improve the finish time. Nonetheless, to our knowledge, there are few previous longitudinal studies on the running pace during the 400-m hurdles event.

Fast sprinting speed (short split time) is determined by high step frequency (SF) and/or high step length (SL); therefore, these two step characteristics are often used to evaluate why faster sprinters can run [13,14] and why sprinters can enhance running speed highly [15–17]. Unfortunately, even cross-sectional studies have not reported the detailed step characteristics of faster 400-m hurdlers in the first and latter halves of the race. The multiple single-subject approach longitudinally shows that SF and/or SL reliance is based on individual changes to maintain high performance in a 100-m race [8] and long jump [18]. Hence, high SF and/or high SL would achieve shortened split times and thereby a shortened finish time in the 400-m hurdles based on individual changes.

The influence of SL and SR on the 100-m finish time is individually different among various sprinters and the influence is not affected by the competitive level [8]. In contrast to the flat sprints, in the hurdles race, hurdlers are not often able to run with an optimal combination between SL and SF to achieve the highest running speed [19], which is externally caused by the fixed inter-hurdle distance [20]. Therefore, if no difference of SL and/or SR is obtained between different competitive-level hurdlers, effect of sensitivities of running pace and the step characteristics on the finish time would be different between different competitive-level hurdlers.

Both cross-sectional and longitudinal approaches would be helpful for individual athletes when determining the target race pace and step characteristics in the 400-m hurdles. Therefore, the purpose of this study was to clarify the running pace and step characteristics between different competitive-level 400-m hurdlers cross-sectionally and clarify whether the hurdlers are individually more reliant on the first-half and/or latter-half pace during the race and more reliant on SF and/or SL in each half of the race. We had three hypotheses: First, do faster 400-m hurdlers run particularly faster in the latter half of the race? Second, are the finish times of hurdlers individually influenced by their race pace and step characteristics? Third, do these influences individually differ between world-class and national-level hurdlers?

## Materials and methods

### Athletes

Because our data set was obtained from publicly-available Internet broadcasts, we did not obtain informed consent.

The sample comprised 13 male world-class and 14 male national-level 400-m hurdlers (best record: 47.71 ± 0.44 s and 49.28 ± 0.41 s, respectively). They were ranked in the top 20 more than three times from 2010 to 2014 in the World and Japan, respectively. This study was conducted after obtaining approval from the Research Ethics Committee involving Living Human Subjects at Ritsumeikan University (BKC-human-2016-053).

### Design

This study was laid out as an observational research design. We obtained the movie data during competitive races from publicly available television and internet broadcasts [8,21-23]. We conducted cross-sectional and longitudinal research approaches to answer the research questions.

### Methodology

The competitions involved Olympic, International Association of Athletics Federation championships, Grand Prix series competitions, national championships, and local competitions in Japan (see, Table S1). In this study, races in which the individual hurdler was likely to run with maximal effort through the finish line were analysed. We disregarded individual races in which the hurdler clearly eased off before the finish line from the analysis. A total of 16.8 ± 4.2 races for each world-class hurdler and 20.0 ± 6.0 races for each national-level hurdler were analysed by publicly available Internet broadcasts [8,21–23]; so the total number of the analysed runs was 499. The research procedures complied with the companies’ terms and conditions for all websites and broadcasts used in this study.

We used the finish time as the official race time. We determined the split-times for the first and latter halves of the race at the instant of touchdown for the leading leg after clearing the fifth hurdle (hereafter, fifth touchdown).

We counted the total number of steps involved in clearing ten hurdles in the race until the first step over the finish line. During the first half of the race, we calculated the mean SL by dividing 185 m (from the start line to the position at the fifth hurdle) by the number of steps until the fifth touchdown and the mean SF by dividing the numbers of steps by the duration from the instant of the gunfire to the instant of the fifth touchdown. During the latter half of the race, we calculated the mean SL by dividing 215 m (from the position at the fifth hurdle to the finish line) by the number of steps to the finish time, and the mean SF by dividing the number of steps by the duration from the fifth touchdown to the finish line. Because most of the hurdlers did not complete a step exactly at 400 m, we calculated the last step from just before the finish line to the finish line (*Step*__*last*__) for each sprinter as follows:

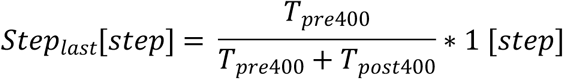

*T*_*pre*400_ is the duration from the touchdown of the step just before the finish line to the finish line. *T*_*post*400_ is the duration from the finish line to the touchdown of the step just after the finish line.

### Statistical Analysis

We present all parameters mean ± SD. We checked all datasets for normality and homogeneity of variance. To assess the differences in mean values and SDs of the finish times, split times, SLs, and SFs between the world-class and national-level hurdlers, we used unpaired T-tests. To assess the cross-sectional relationships between two parameters, we used Pearson’s correlations coefficients.

In the multiple single-subject approach, we analysed each hurdler individually [8,17]. We natural log-transformed finish time, split time, mean SF, and mean SL before analysis to normalize distributions and stabilize variance. To determine any first- or latter-half speed reliance for an individual hurdler, we derived the 90% confidence interval (CI) for the difference between first-half split time vs. finish time and latter-half split time vs. finish time relationships (hereafter, first-half speed reliant index) using a criterion nonparametric bootstrapping technique (Matlab R2017a, Marthworks Inc., Natick, MA). Similarly, we calculated the 90% CI for the difference between the SL vs. split time and SF vs. split time relationships (hereafter, SL reliant index) in the first and latter halves of each race. We used Pearson’s (*r*) correlations to calculate the relationships between the differences for each resample. We employed a bootstrapping technique to provide 10,000 resamples of the natural log transformed finish time, split time, mean SL, and mean SF values. We identified hurdlers as being first-half speed reliant or SL reliant if the mean correlation difference was positive with the lower limit of the 90% CI ≥ −0.1; in contrast, we identified them as being latter-half speed reliant or SF reliant if the mean correlation difference was negative with the lower limit of the 90% CI ≤ 0.1. Salo et al. [8] present this data analysis in detail.

To assess the differences in the first-half speed reliant and SL reliant indices between world-class and national-level athletes, we used non-parametric Mann-Whitney’s *U* tests. Eta-squired was calculated for assessing the sample size in Mann-Whitney’s *U* tests.

We set statistical significance at *P* < 0.05.

## Results

### Cross-sectional research approach

Mean values of first- and latter-half split times and finish times in all trials of the world-class hurdlers were significantly shorter than those of the top national-level hurdlers (Table 1). Greater mean SF contributed significantly to the shorter mean first-half split times of the world-class hurdlers compared to those of the national-level athletes. In contrast, no significant differences in the SDs for finish time, split time, and mean SF in all trials were obtained between the two different groups. However, the SD for the latter-half mean SL of the world-class hurdlers was significantly larger than that of national-level hurdlers. As with the other parameters, the personal best times in the 400-m race of the world-class hurdlers were significantly shorter than those of the national-level hurdlers (45.74 ± 1.01 s vs. 47.42 ± 0.65 s, *P* < 0.01); in contrast, no significant difference in body height was observed between the two different groups (1.84 ± 0.07 m vs. 1.79 ± 0.05 m, n.s.).

**Table 1.** Cross-sectional mean value and SD in all trials of spatiotemporal parameters of world-class and national-level hurdlers (mean ± SD)

|  | World-class hurdlers | National- level hurdlers |
| --- | --- | --- |
| **Mean value** |  |  |
| **T<sub>total</sub> [s]** | $48.85 \pm 0.33$ * | $50.43 \pm 0.44$ |
| **T<sub>first</sub> [s]** | $21.41 \pm 0.29$ * | $22.18 \pm 0.24$ |
| **T<sub>latter</sub> [s]** | $27.43 \pm 0.23$ * | $28.23 \pm 0.37$ |
| **SR<sub>total</sub> [Hz]** | $3.58 \pm 0.14$ | $3.50 \pm 0.10$ |
| **SR<sub>first</sub> [Hz]** | $3.74 \pm 0.12$ * | $3.60 \pm 0.11$ |
| **SR<sub>latter</sub> [Hz]** | $3.47 \pm 0.16$ | $3.43 \pm 0.11$ |
| **SL<sub>total</sub> [m]** | $2.29 \pm 0.09$ | $2.27 \pm 0.07$ |
| **SL<sub>first</sub> [m]** | $2.32 \pm 0.09$ | $2.32 \pm 0.08$ |
| **SL<sub>latter</sub> [m]** | $2.27 \pm 0.10$ | $2.23 \pm 0.08$ |
| **SD** |  |  |
| **T<sub>total</sub> [s]** | $0.64 \pm 0.13$ | $0.62 \pm 0.12$ |
| **T<sub>first</sub> [s]** | $0.31 \pm 0.08$ | $0.32 \pm 0.07$ |
| **T<sub>latter</sub> [s]** | $0.54 \pm 0.10$ | $0.52 \pm 0.11$ |
| **SR<sub>total</sub> [Hz]** | $0.05 \pm 0.02$ | $0.04 \pm 0.01$ |
| **SR<sub>first</sub> [Hz]** | $0.07 \pm 0.06$ | $0.05 \pm 0.01$ |
| **SR<sub>latter</sub> [Hz]** | $0.06 \pm 0.02$ | $0.06 \pm 0.01$ |
| **SL<sub>total</sub> [m]** | 0.02 ± 0.02 | 0.02 ± 0.01 |
| **SL<sub>first</sub> [m]** | 0.03 ± 0.05 | 0.02 ± 0.02 |
| **SL<sub>latter</sub> [m]** | 0.03 ± 0.01 * | 0.02 ± 0.01 |

**Note.** T<sub>total</sub> = finish time; T<sub>first</sub> = split time during the first-half phase; T<sub>latter</sub> = split time during the latter-half phase; SR<sub>first</sub> = mean step rate during the first-half phase; SL<sub>first</sub> = mean step length during the first-half phase; SR<sub>latter</sub> = mean step rate during the latter-half phase; SL<sub>latter</sub> = mean step length during the latter-half phase. \*Significant difference from national-level hurdlers (P < 0.05).

Positive relationships of finish time were highly obtained between first- and latter-half split times (Fig 1). In contrast, no significant relationships of split-time were obtained between mean SF and mean SL in either the first or latter half of the race. The finish time of the 400-m hurdles was significantly related to personal records in the 400-m race (r = 0.798, *P* < 0.01).

**Fig 1.**
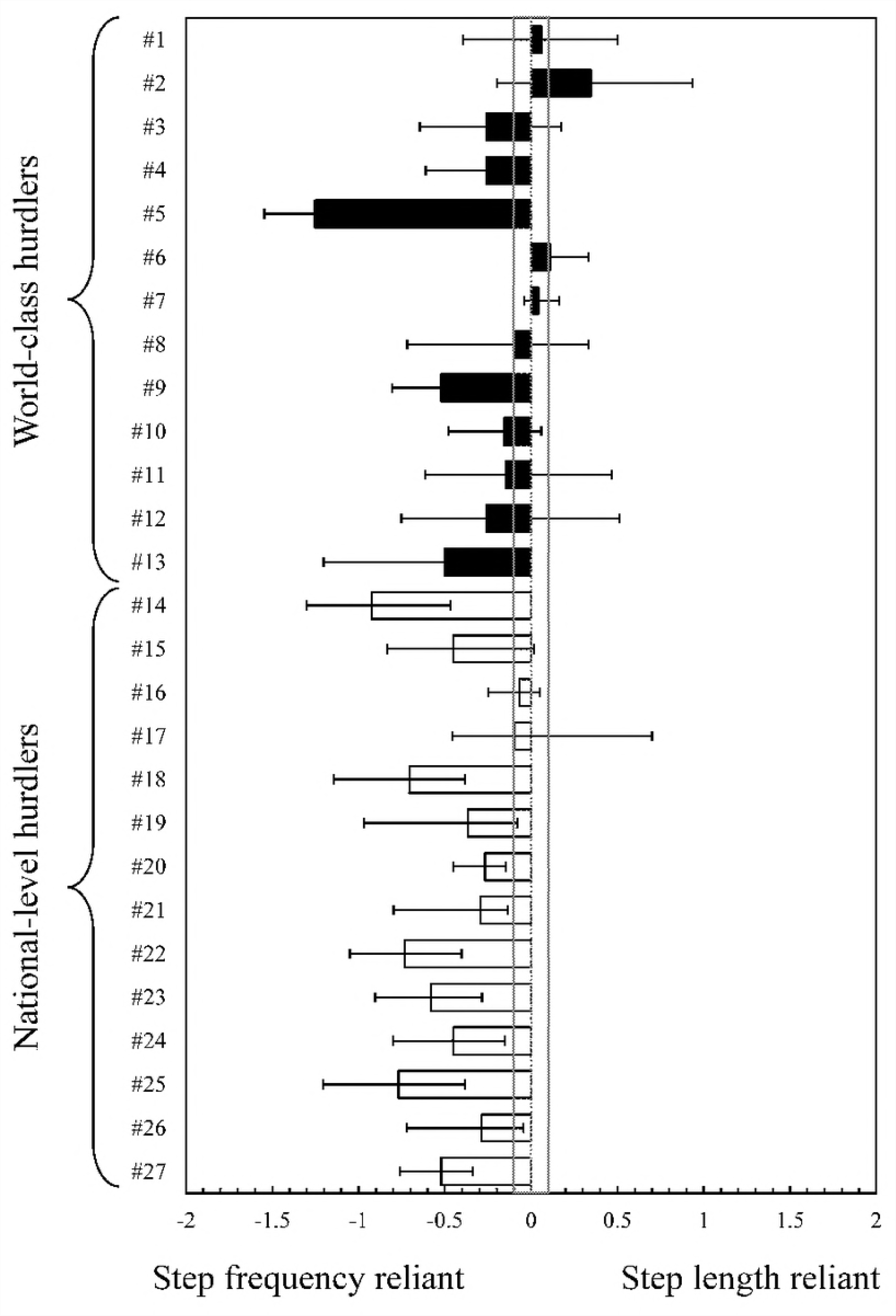
Scatter plots for finish times, split times, and step characteristics by cross-sectional design. Filled circles denote data of world-class hurdlers and open circles denote data of national-level hurdlers.

### Longitudinal research approach

Correlation coefficients between each of the parameters within single subjects are presented in Table 2. The finish time-split time correlations for 26 of the 27 hurdlers were positive and the magnitudes ranged from 0.10 to 0.97. A greater number of correlations over 0.70 were obtained in the latter half of the race (26/27) compared to the first-half (5/27). In both the first and latter halves of the race, the split time-mean SL correlations for most of the hurdlers were negative and the magnitudes ranged from 0.08 to 1.00. A greater number of correlations over 0.70 were obtained in split time-mean SF compared to split time-mean SL in both the first (19/27 vs. 2/27) and latter halves (22/27 vs. 4/27) of the race.

**Table 2.** Within-subject correlation coefficients among spatiotemporal parameters.

| Participant | Split time |  | First-half phase |  | Latter-half phase |  |
| --- | --- | --- | --- | --- | --- | --- |
| | $T_{\text{total}}-T_{\text{first}}$ | $T_{\text{total}}-T_{\text{latter}}$ | $T_{\text{first}}-SR_{\text{first}}$ | $T_{\text{first}}-SL_{\text{first}}$ | $T_{\text{latter}}-SR_{\text{latter}}$ | $T_{\text{latter}}-SL_{\text{latter}}$ |
| #1 | 0.52 | 0.84 | -1.00 | 0.00 | -0.41 | -0.47 |
| #2 | 0.25 | 0.88 | -1.00 | 0.00 | -0.40 | -0.75 |
| #3 | 0.80 | 0.90 | -0.45 | -0.49 | -0.73 | -0.47 |
| #4 | 0.48 | 0.88 | -0.58 | -0.01 | -0.81 | -0.55 |
| #5 | 0.76 | 0.97 | -1.00 | 0.00 | -0.99 | 0.26 |
| #6 | 0.45 | 0.76 | -0.98 | -0.04 | -0.74 | -0.85 |
| #7 | 0.53 | 0.93 | -0.96 | -0.14 | -0.82 | -0.86 |
| #8 | 0.85 | 0.78 | -0.63 | -0.25 | -0.57 | -0.48 |
| #9 | 0.34 | 0.87 | -0.90 | 0.10 | -0.95 | -0.43 |
| #10 | 0.85 | 0.87 | -0.08 | -0.20 | -0.79 | -0.63 |
| #11 | 0.54 | 0.82 | -1.00 | 0.00 | -0.73 | -0.58 |
| #12 | 0.59 | 0.97 | -0.72 | 0.09 | -0.62 | -0.36 |
| #13 | -0.10 | 0.87 | -1.00 | 0.00 | -0.94 | -0.44 |
| #14 | 0.81 | 0.73 | -0.91 | -0.24 | -0.78 | 0.14 |
| #15 | 0.48 | 0.81 | 0.18 | -0.71 | -0.84 | -0.39 |
| #16 | 0.67 | 0.91 | -1.00 | 0.00 | -0.78 | -0.71 |
| #17 | 0.61 | 0.80 | -0.94 | -0.49 | -0.64 | -0.54 |
| **#18** | 0.68 | 0.87 | -0.95 | -0.14 | -0.89 | -0.18 |
| **#19** | 0.51 | 0.84 | -0.97 | -0.41 | -0.98 | -0.61 |
| **#20** | 0.55 | 0.92 | -0.98 | 0.41 | -0.96 | -0.69 |
| **#21** | 0.51 | 0.92 | -0.28 | -0.68 | -0.90 | -0.61 |
| **#22** | 0.46 | 0.68 | -0.64 | -0.26 | -0.89 | -0.16 |
| **#23** | 0.66 | 0.91 | -0.98 | 0.18 | -0.87 | -0.29 |
| **#24** | 0.46 | 0.87 | -0.98 | 0.02 | -0.82 | -0.37 |
| **#25** | 0.58 | 0.87 | -0.54 | -0.70 | -0.89 | -0.12 |
| **#26** | 0.24 | 0.84 | -0.89 | 0.05 | -0.78 | -0.49 |
| **#27** | 0.61 | 0.87 | -1.00 | 0.00 | -0.90 | -0.37 |

Note. $T_{\text{total}}$ = finish time; $T_{\text{first}}$ = split time during the first-half phase; $T_{\text{latter}}$ = split time during the latter-half phase; $SR_{\text{first}}$ = mean step rate during the first-half phase; $SL_{\text{first}}$ = mean step length during the first-half phase; $SR_{\text{latter}}$ = mean step rate during the latter-half phase; $SL_{\text{latter}}$ = mean step length during the latter-half phase.

More than half (14/27) of the athletes were identified as being latter-half speed reliant; in contrast, no athletes demonstrated first-half speed reliance and the 13 remaining athletes did not favour either characteristic (Fig 2). In the first-half of the race, 19 of the 27 athletes were identified as being SF reliant, only one athlete demonstrated SL reliance, and the remaining seven athletes did not favour either characteristic (Fig 3). In the latter half of the race, 12 of the 27 athletes were identified as being SF reliant, no athletes demonstrated SL reliance, and the remaining 15 athletes did not favour either characteristic (Fig 4).

**Fig 2.**
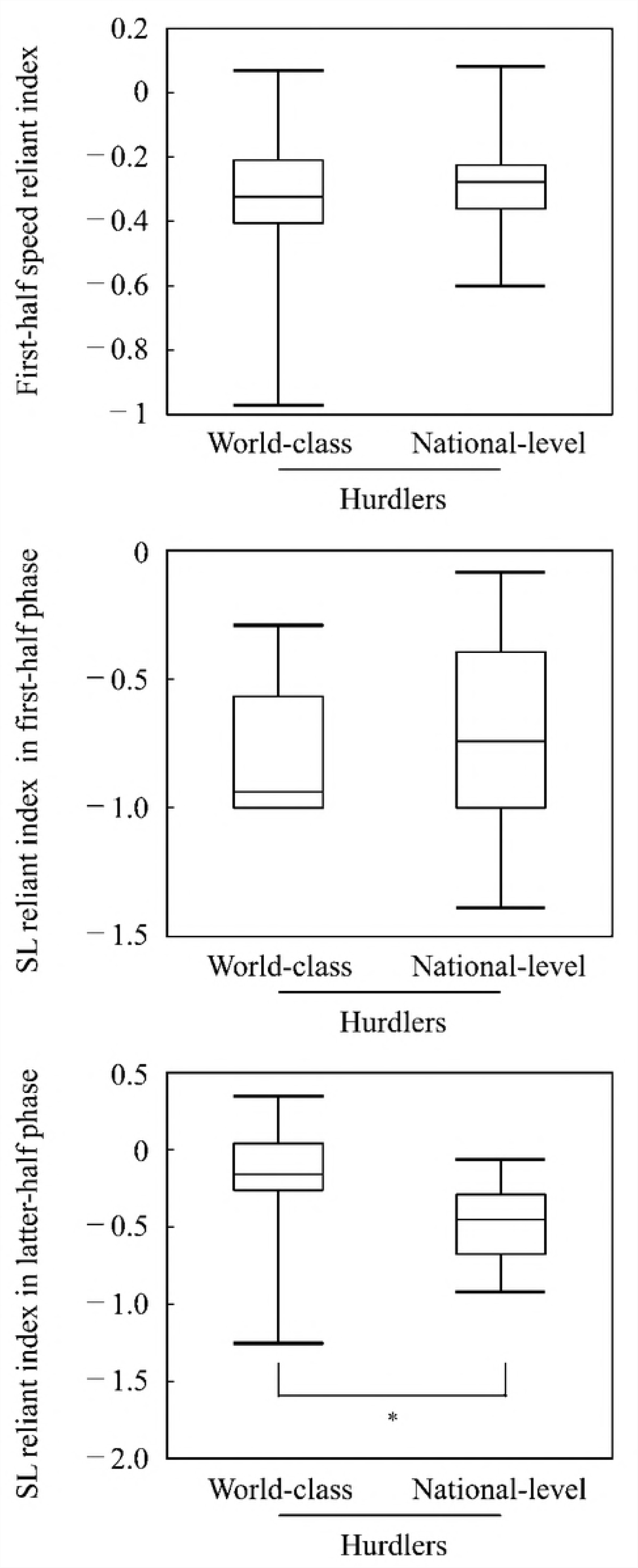
The first-half speed reliant index with 90% confidence interval for each hurdler (#1 to #27). The area ±0.1 from zero in the middle demonstrates the trivial (nonreliant) effect. The black bars denote world-class hurdlers and the white bars denote national-level hurdlers.

**Fig 3.**
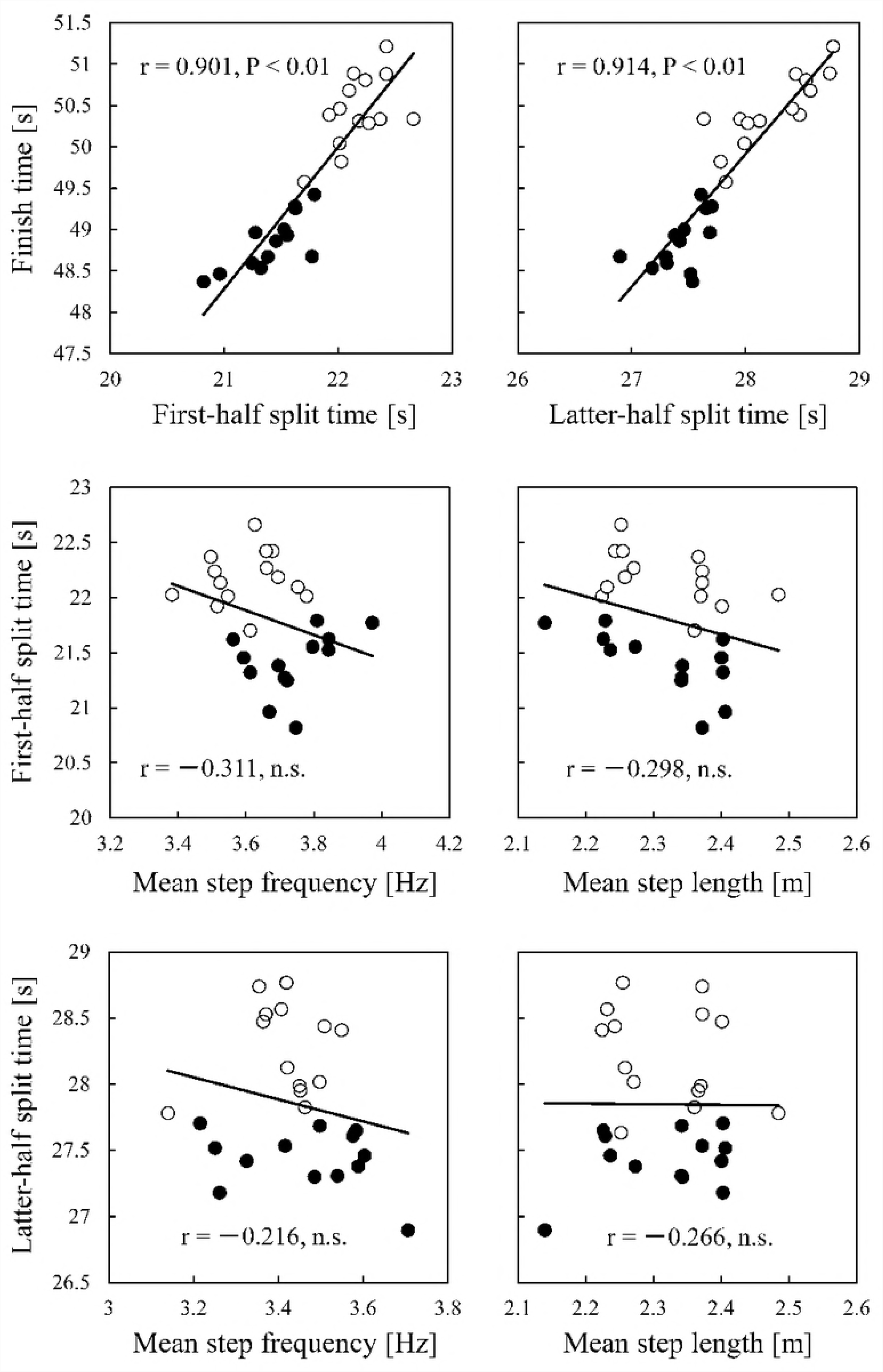
The step length reliant index during the first-half of the race with 90% confidence interval for each hurdler (#1 to #27). The area ±0.1 from zero in the middle demonstrates the trivial (nonreliant) effect. Black bars denote world-class hurdlers and white bars denote national-level hurdlers.

**Fig 4.**
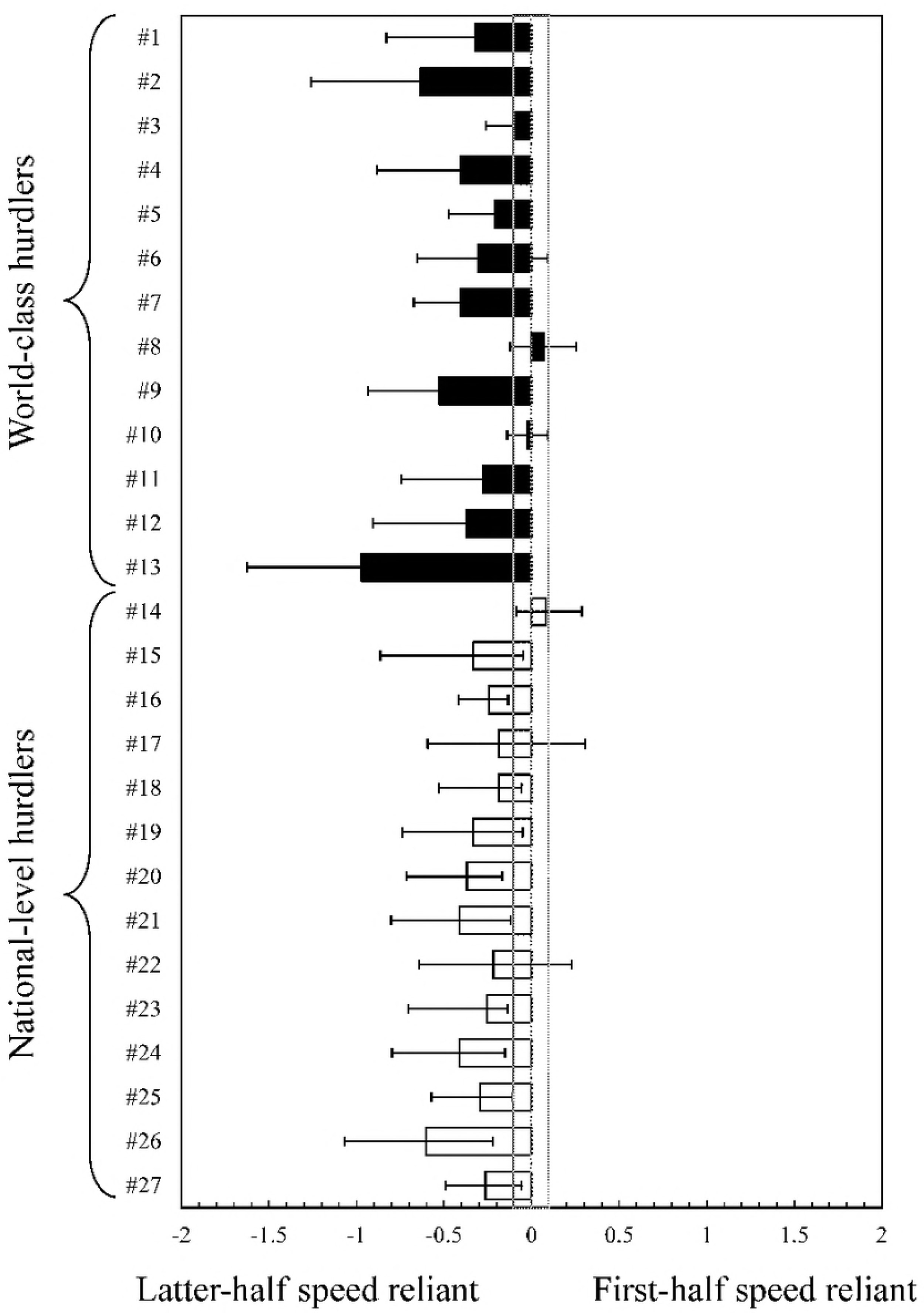
The step length reliant index during the latter half of the race with 90% confidence interval for each hurdler (#1 to #27). The area ±0.1 from zero in the middle demonstrates the trivial (nonreliant) effect. Black bars denote world-class hurdlers and white bars denote national-level hurdlers.

No significant differences in the first-half speed reliant (η^2^= 0.01) and the SL reliant indices (η^2^ = 0.00) were observed between world-class and national-level hurdlers (Fig 5). In contrast, the SL reliant index of the world-class hurdlers was significantly larger than that of national-level athletes (η^2^ = 0.22).

**Fig 5.**
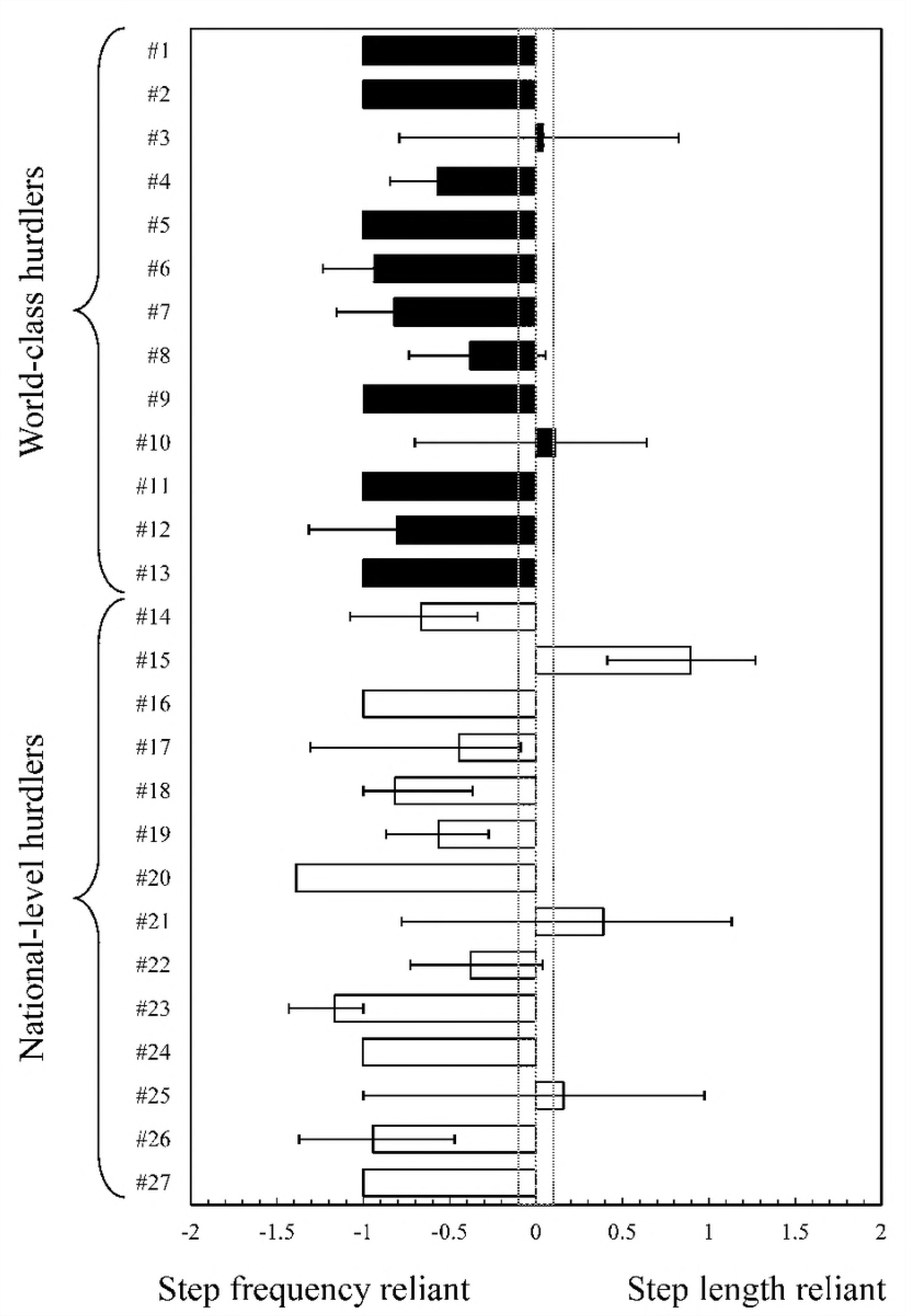
Box plot of the first-half speed reliant index and step length (SL) reliant index in first- and latter-half phases, comparing world-class and national-level hurdlers. ^*^indicates *P* < 0.05.

## Discussion

This study was the first to clarify the running paces and step characteristics of 400-m hurdlers using both cross-sectional and longitudinal research approaches. Our main findings were as follows: first, even if the cross-sectional approach showed the extreme importance of the split time during both first- and latter-halves of the race, the multiple single-subject approach longitudinally showed that the latter-half split time was important for all hurdlers to better their finish times. Second, the faster hurdlers’ step characteristics were not different between the slower hurdlers’ those throughout the races; in contrast, mean SF in the latter half of the race was important for all the hurdlers to decrease their split-times. Third, the influence of mean SL on the split time in the latter half of the race was larger for the world-class hurdlers than for the national-level hurdlers. Therefore, the correlations in the cross-sectional research approach do not directly indicate causation in the 400-m hurdles, and the individualization principle for elite hurdler training can be considered essential.

To our knowledge, there have been few previous studies on race pace and step characteristics in the 400-m hurdles. One previous study showed that the correlation coefficients of the finish time between first- and latter-half split times were 0.359 to 0.737 [12]; in contrast, those obtained in this study were clearly higher (0.900 and 0.913). This difference may be because the spatiotemporal data in this study were averaged from many competitive races for each hurdler (17.9 ± 4.3 races for world-class hurdlers and 21.2 ± 5.9 races for national-level hurdlers) so the acute daily effects of the sudden wind speed [24] and different race pace can be considered smaller compared to the analysis of only one race. In fact, the within-subject SDs of the first- and latter-half split times ranged widely over many competitive races (e.g., latter-half split time of world-class hurdlers: 0.54 ± 0.10 s). This finding suggests that analysing only one race for each hurdler cannot reflect the athlete’s normal performance; so, our cross-sectional methodology, which used the mean value for various races, can be considered as more accurate than that in a previous study [12].

A strong positive relationship between finish times and latter-half split times was observed in the cross-sectional research approach, demonstrating that our first hypothesis was accepted. Moreover, great number of the finish time-split time correlations over 0.70 were obtained in the latter half of the race. Interestingly, there were no hurdlers who sacrificed the latter-half split time in favour of a faster first-half split time (first-half speed reliant hurdlers: 0/27), suggesting our second hypothesis was partially not accepted. These findings demonstrate that a shorter split-time in the latter half of the race is key for all hurdlers to achieve better personal finish times. Good split-times in the latter half of a long sprint race are determined by several factors: small or optimal energy consumption in the first-half of the race [5], increased utilization of glycolytic and aerobic systems [6], good technique while clearing the hurdles [25,26], and external environment [24]. Unfortunately, we cannot determine the factors in detail because of our methodology; however, these factors might affect the high performance of individuals in the latter half of the race.

The multiple single-subject approach clarified that, for most hurdlers, the short split-time was not related to high mean SL, but was associated with high mean SF during both the first and latter halves of the race. At high sprinting speeds in short sprint running, increases in step frequency have a greater effect on increases in sprinting speed [14,16,27]. This finding corresponded with our results; thus, split time can be considered as more sensitive to mean SF rather than mean SL in a 400-m hurdles race. On the other hand, in a 400-m race, the differences in running speed among different level male sprinters are not caused by the differences in mean SF, but rather by the differences in mean SL throughout the whole race [11]. In contrast, our cross-sectional approach results did not support this finding: no relationships were observed between split times and SL during both the first and latter halves of the race. This finding may be due to the narrower range of finish times among our subjects compared to those in a previous study [11], which does not have as significant an effect on the relationship. Moreover, this difference might be due to the fact that some hurdlers could not run with an optimal combination between mean SL and mean SF to achieve the highest running speed [19] because they were forced to run between hurdles in a fixed step rhythm [20].

In the latter half of the race, although no significant difference in mean SL was observed between the two hurdler groups, the within-subject SD of the SL of the world-class hurdlers was significantly greater than that of the national-level hurdlers. As abovementioned, hurdles are not often able to run with an optimal SL for high running speed because of the fixed inter-hurdle distance [20]. The 400-m personal records of the world-class hurdlers were greater than those of the national-level hurdlers, corresponding with the findings in a previous study [28]. Therefore, the SL of the world-class hurdlers might not be more optimal for fast running compared with that of the national-level hurdlers in the latter half of the race. This might lead the SL index of the world-class hurdlers to be higher than that of the national-level hurdlers. Thus, our third hypothesis was accepted.

From the results in this study, we can provide the following information as practical applications for the 400-m hurdlers. First, when developing training programs and pace strategies, all hurdlers should prioritize improving step characteristics and running speed in the latter half of the 400-m hurdles rather than in the first half of the race. Second, improving performance in the first half of the race should be considered based on the individualization principle for training.

We have a limitation in this study: we could not measure the parameters to assess biochemical energy expenditure. Long sprint running is performed by the glycolytic system with chemical energy supplies involving adenosine triphosphate and phosphocreatine, as well as aerobic systems [2–5,29]. This biochemical energy is finally converted to mechanical energy and various biomechanical parameters [5,7], leading us to answer our question with a detailed explanation. However, during competition, it is very difficult to obtain these parameters. Further research is needed to answer our questions using parameters for biochemical energy expenditure.

## Conclusion

In conclusion, although the cross-sectional approach showed strong correlations between the finish time and performance in the first and latter halves of the race, the multiple single-subject approach showed latter-half performance is more essential for all hurdlers. Therefore, the important findings regarding high performance in a cross-sectional study approach do not always correspond with those in a longitudinal approach.

